# StabMap: Mosaic single cell data integration using non-overlapping features

**DOI:** 10.1101/2022.02.24.481823

**Authors:** Shila Ghazanfar, Carolina Guibentif, John C. Marioni

## Abstract

Currently available single cell -omics technologies capture many unique features with different biological information content. Data integration aims to place cells, captured with different technologies, onto a common embedding to facilitate downstream analytical tasks. Current horizontal data integration techniques use a set of common features, thereby ignoring non-overlapping features and losing information. Here we introduce StabMap, a mosaic data integration technique that stabilises mapping of single cell data by exploiting the non-overlapping features. StabMap is a flexible approach that first infers a mosaic data topology, then projects all cells onto supervised or unsupervised reference coordinates by traversing shortest paths along the topology. We show that StabMap performs well in various simulation contexts, facilitates disjoint mosaic data integration, and enables the use of novel spatial gene expression features for mapping dissociated single cell data onto a spatial transcriptomic reference.

## INTRODUCTION

Large-scale efforts to build transcriptional maps of tissues at cellular resolution have revealed many biological insights and provided reference maps that can be used to further interrogate biological systems^1,2^. Simultaneous technological advances have led to the generation of datasets that capture multiple distinct types of molecular information, for example, CITE-seq captures RNA expression and cell surface protein abundance^3^, and 10X Genomics Multiome captures RNA expression alongside DNA fragments associated with regions of open chromatin^4^. Consequently, data integration has emerged as a key challenge for consolidating and profiting from such rich resources^5^, with the task of integrating diverse molecular assays being known as ‘mosaic data integration’^6^. At present, many methods for mosaic data integration are typically limited to using the set of overlapping features between modalities^7,8^.

However, as the number and complexity of single cell datasets increase, there is a growing need to develop techniques specifically designed to perform mosaic data integration^9,10^. Some existing approaches designed to tackle this problem include UINMF^11^, which introduces a latent metagene matrix in the factorisation problem, and MultiMAP^12^, a graph-based method that assumes a uniform distribution of cells across a latent manifold structure fitted using an optimisation approach. A critical limitation of both approaches, however, is the requirement that all datasets contain some features that are shared across all datasets, resulting in analysts needing to compromise on input datasets, or making the ‘central dogma assumption’, i.e. matching features between different -omics modalities based on corresponding DNA-RNA-protein sequences. Moreover, while MultiMAP includes a tuning parameter to prioritise certain datasets, neither approach offers a supervised mode that takes into account *a priori* cell labels.

In this paper, we introduce StabMap, a data integration technique designed specifically for mosaic data integration tasks. StabMap projects all cells onto supervised or unsupervised reference coordinates utilising all available features regardless of overlap with other datasets, instead relying on traversal along the mosaic data topology. By using multiple simulation scenarios and by exploring spatially resolved transcriptomic data, we show that StabMap performs well, in particular in the presence of very few overlapping features. Additionally, we demonstrate StabMap’s novel ability to perform disjoint mosaic data integration, and reveal new biological insights into the role of Brachyury in early mouse organogenesis.

## RESULTS

### StabMap: stabilised mapping for mosaic single cell data integration

The input to StabMap is a set of single cell data matrices, and an optional set of discrete cell labels. From this data structure StabMap extracts the mosaic data topology (MDT), a network with nodes corresponding to each given dataset, and edges between nodes, weighted by the absolute number of shared features between the datasets (Figure 1A). StabMap only requires that the MDT be a connected network, i.e. there be a way to draw a path from every node to every other node. For the selected reference dataset, R, a supervised (Linear Discriminant Analysis (LDA), if labels provided) or unsupervised (PCA) dimensionality reduction algorithm is employed, generating a features loading matrix for the discriminants or components. This is performed using all features available for the reference dataset. Then, for each non-reference dataset, D, the shortest path is identified between R and D along the MDT. If there is a direct link between R and D, a multivariable linear model is fitted to estimate the PC and/or LD scores, with predictor variables corresponding to the shared features between datasets R and D. If there is no direct link between R and D, StabMap will construct a sequence of mappings between features traversing the shortest path between R and D along the MDT by iteratively predicting the scores of the reference dataset (Figure 1B, Methods). In the case where multiple datasets are considered as reference datasets, the above process is repeated and the mappings are then concatenated to form a single low-dimensional matrix (Methods). The resulting StabMap embedding can be employed for further downstream analysis tasks, including batch correction, joint visualisation, supervised and unsupervised machine learning tasks, differential abundance testing, and testing for and characterising developmental trajectories.

**Figure 1.**
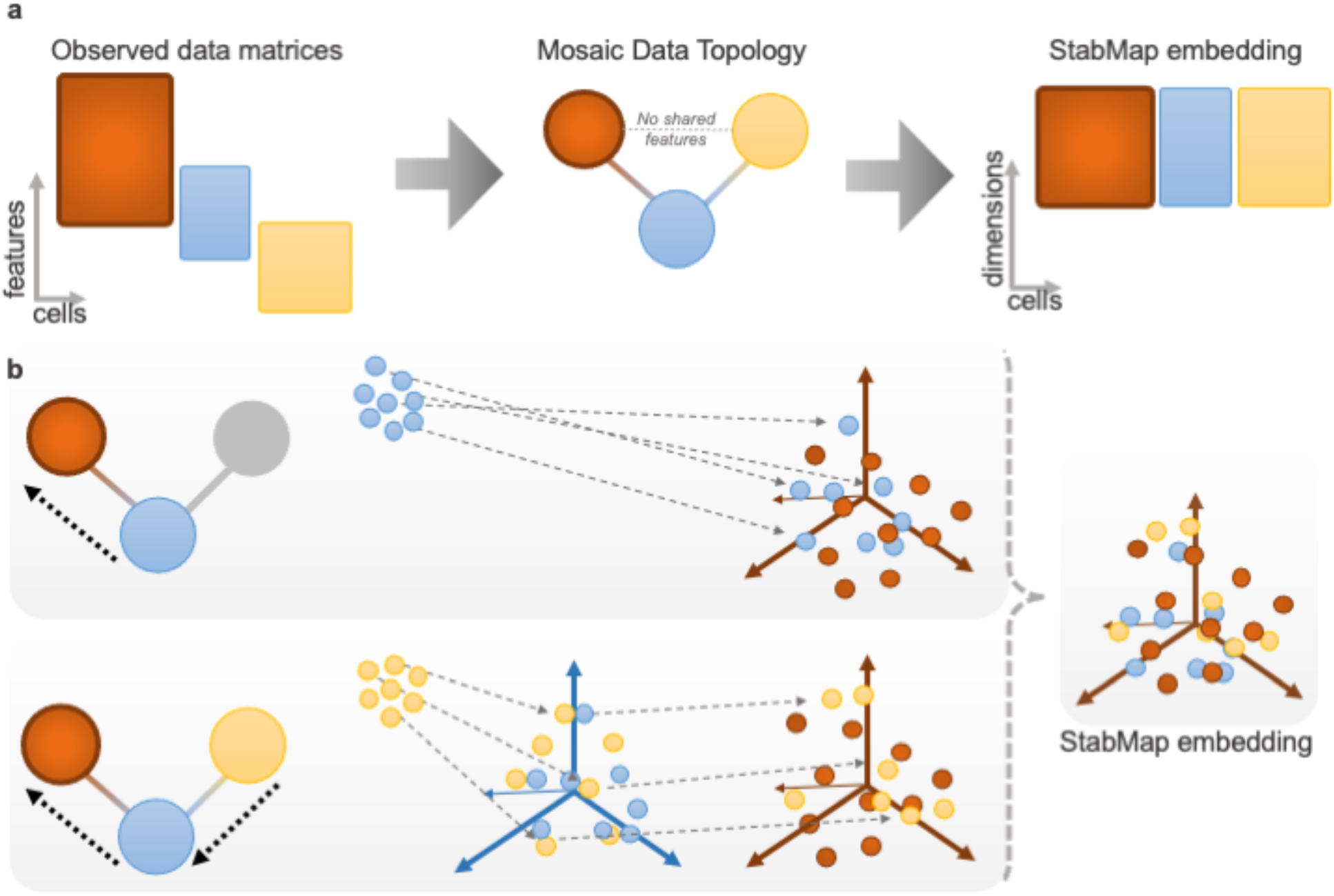
StabMap method overview. **a**. Example mosaic data integration displaying observed data matrices with varying overlap of features among the datasets. Datasets are summarised using the mosaic data topology (MDT). Cells are then projected onto the common StabMap embedding across all cells. **b**. Cells from all datasets are projected onto the reference space (dark red) by traversing the shortest paths along the MDT. Blue cells are projected directly onto the reference space, whereas yellow cells are first projected onto the space defined by the blue cells, followed by projection to the dark red space. All cells are then combined to yield the common StabMap embedding.

By performing mosaic data integration using traversal along the mosaic data topology, and not relying on the features common to all datasets, StabMap unlocks the ability to perform disjoint mosaic data integration, i.e. integrating data where the intersection of features measured for all datasets is empty.

### StabMap preserves cell-cell relationships in multiomics data

To investigate the performance of StabMap, we first constructed a simulation scenario using multiomics single cell data, where chromatin accessibility and mRNA expression were measured in each of ∼36,000 peripheral blood mononuclear cells (PBMCs)^13^. Using these data, we computationally created two single-cell datasets - one containing only the mRNA measurements and the other only the chromatin accessibility measurements - and assumed that the problem of interest was to combine these two datasets onto a common scaffold. We used all highly variable genes from the RNA modality, and all highly variable peaks from the ATAC modality, and considered the peaks associated with promoter regions of genes as common features (Figure 2A).

**Figure 2.**
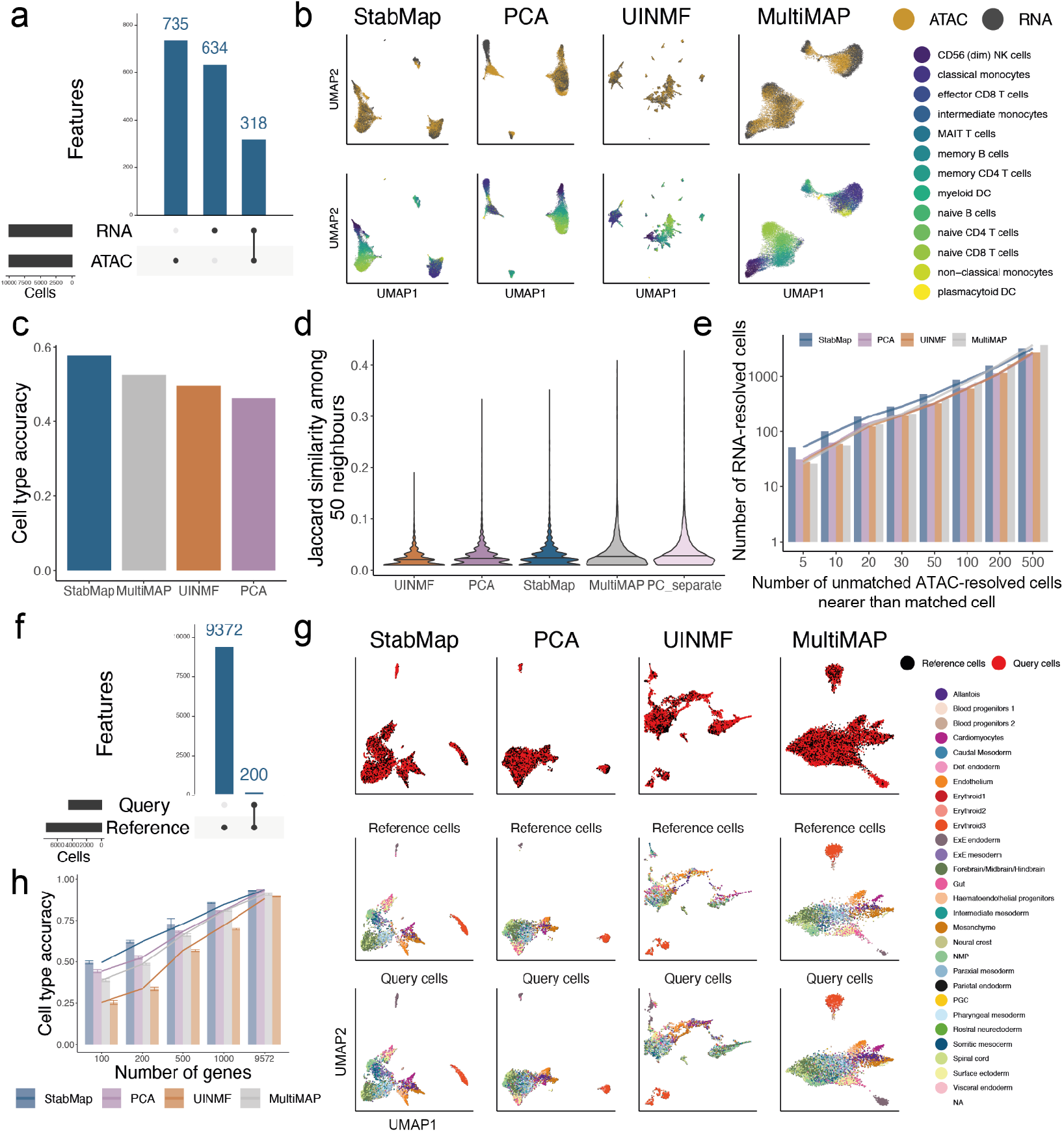
Mosaic data integration simulations using PBMC Multiome and Mouse Gastrulation Atlas data. **a**. UpSet plot of features shared between simulated RNA and ATAC modalities. ATAC peaks in promoter regions of genes are aligned with the genes in the RNA modality. **b**. UMAP representations of RNA and ATAC modality cells for StabMap (first column), PCA, UINMF and MultiMAP (last column), coloured by simulated modality (top row) and by cell type (bottom row). **c**. Barplot of cell type classification accuracy predicting ATAC-resolved cell types using RNA-resolved cells as training data. **d**. Violin plots displaying Jaccard similarity among 50 neighbours for cells in each modality, where a higher value indicates a better preservation of neighbourhood structure. **e**. Barplot displaying the cumulative number of RNA-resolved cells, grouped by the number of unmatched ATAC-resolved cells found to be nearer than the matched ATAC-resolved cell. A higher overall curve indicates more cells are closer to their true neighbour. **f**. UpSet plot of features between simulated query and reference datasets for Mouse Gastrulation Atlas data. **g**. UMAP representations of Mouse Gastrulation Atlas data simulation using StabMap, PCA, MultiMAP, and UINMF. First row shows the query cells coloured by cell type, second row shows reference cells coloured by cell type, and the third row shows query cells coloured by cell type. **h**. Barplot displaying the cell type classification accuracy of query cells for various methods, when the query set is restricted to different numbers of genes.

Within this context, we compared StabMap’s performance with i) a naive approach where PCA was applied only to overlapping features, ii) with UINMF and iii) with MultiMAP. In general, we observed reasonable mixing of the RNA- and ATAC-simulated cells with each other across all four computational approaches, as well as distinct separation of cell types (Figure 2B). However, when assessing performance using more quantitative metrics, including the accuracy with which cell types could be predicted (when using the ATAC as the testing set and the RNA as the training set) and the preservation of the distances between cells in the common space, we noted more substantial differences (Methods; Figure 2C-E). Specifically, we observed that while StabMap generally performed well, the other methods (especially the naive PCA implementation and UINMF) had difficulty in accurately predicting cell type (Figure 2C) and in preserving local neighborhood structure (Figure 2E). Taken together, these results suggest that StabMap is well able to perform mosaic data integration.

### StabMap has superior performance when only non-optimal features are available

To further investigate the properties of StabMap, we used scRNA-seq data generated to study mouse gastrulation across entire embryos and at multiple time points^1^ in order to simulate a mosaic data integration task where the reference data contains an assay that captures the full transcriptome (i.e. from scRNA-seq), and the query data contains only a subset of the available gene expression features (e.g. as would be the case for technologies such as seqFISH^14^, MERFISH^15^, qPCR, etc.). We considered the situation where the most informative features are not necessarily known a priori, and split the cells into two datasets, for which one was assumed to contain a small number of genes (n = 100, 200, 500, 1000) randomly selected from among the highly variable genes in the full transcriptome data (Figure 2F; Methods). We compared StabMap with UINMF, MultiMAP and PCA, and visually noted the decrease in structure apparent amongst the query cells in the common embedding for these other methods compared to StabMap (Figure 2G, Supplementary Figure 1). A common task when mapping a query dataset to a reference dataset is to predict the cell types of the query cells. Consequently, we assessed the quality of the data integration task by calculating the K-nearest neighbours cell type classification accuracy (Methods). We identified a much higher accuracy for StabMap, especially when very few features were captured in the simulated query datasets (Figure 2H). Taken together, our results suggest that StabMap is effective at stabilising mapping between datasets even when some of the datasets / modalities contain non-optimal features.

### The robustness of StabMap for disjoint mosaic data integration

Since StabMap relies on the mosaic data topology of the datasets, multiple datasets where some pairs of datasets do not share any features can be embedded into the same StabMap space. This contrasts with existing implementations of PCA, UINMF and MultiMAP, all of which require at least one feature to be shared across all datasets. While this is a major advantage of StabMap, we reasoned that its ability to perform disjoint mosaic data integration would depend heavily on the quality of the input datasets. Consequently, we established how reliably StabMap was able to perform disjoint mosaic data integration with differing levels of information content. Using the 10X Genomics PBMC Multiome data, we randomly split the cells equally into three simulated data types, RNA only, ATAC only, and Multiome (Methods). We intentionally opted to not assign ATAC promoter peak IDs to gene names (i.e. opting to not make the “central dogma assumption”), to replicate the disjoint mosaic data integration task, such that there are no explicitly shared features between the RNA only and ATAC only datasets (Figure 3A-C). We observed that StabMap successfully integrated these three datasets, with cells evenly distributed by data modality, and distinct cell type identities being clearly visible (Figure 3D). Since the most connected node in the MDT is the Multiome dataset, we next queried whether the quality of the StabMap embedding would deteriorate when fewer cells were present in this Multiome dataset. Indeed, we found that when fewer than ∼1,000 cells were allocated to the Multiome dataset, the quality of the StabMap embedding was compromised, with poor local inverse Simpson’s index (LISI)^16^ values relative to modality and cell type (Figure 3E, Supplementary Figure 2). However, when the ‘bridge’ datasets contained more than 1,000 cells we observed highly consistent performance, suggesting that disjoint mosaic integration with StabMap is robust as long as a moderately sized bridge dataset is present.

**Figure 3.**
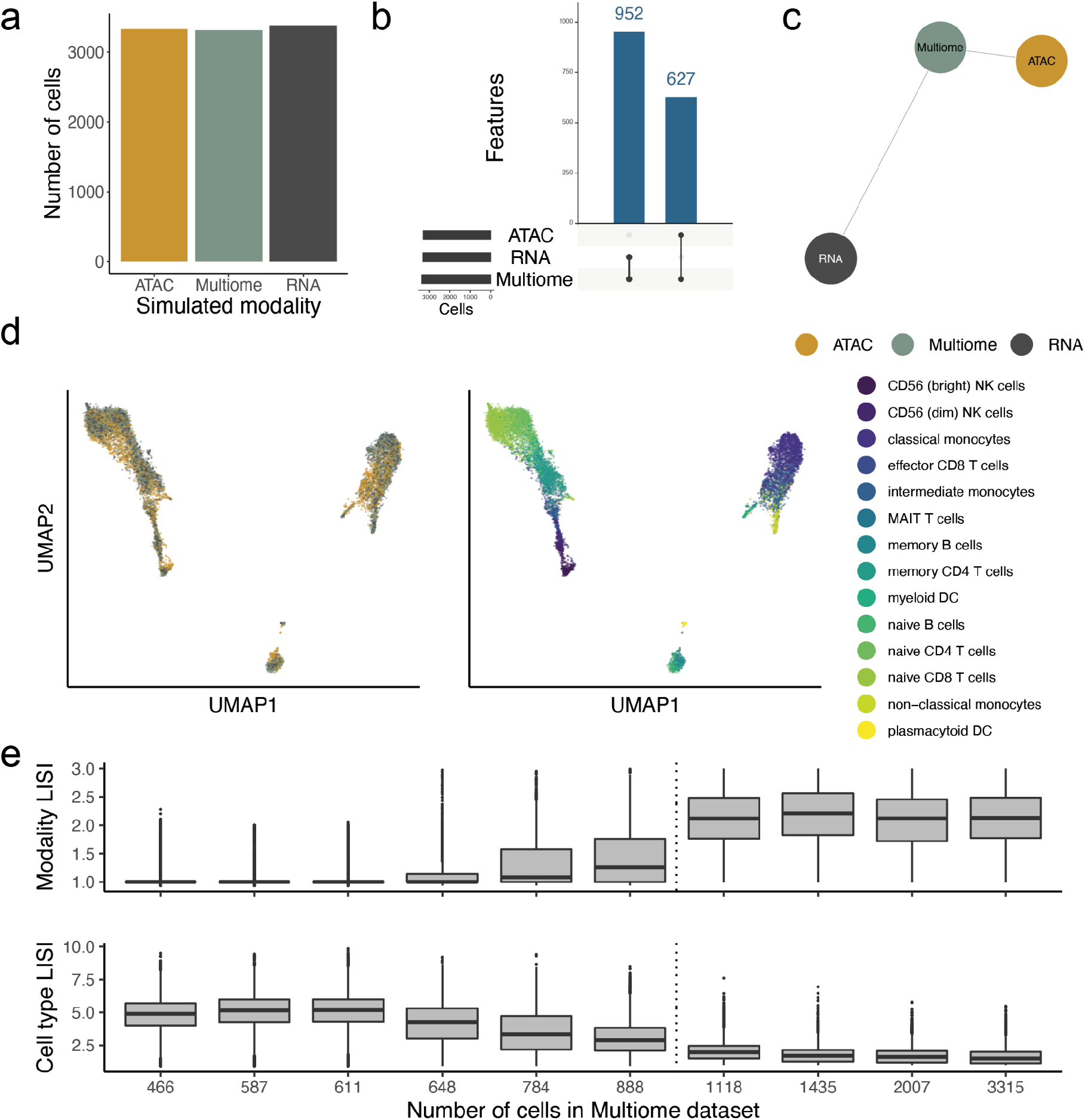
Disjoint mosaic data integration simulation. **a**. Number of multiome cells assigned to each simulated data type. **b**. UpSet plot of overlapping features between simulated data types. **c**. Mosaic data topology of these datasets. **d**. Joint UMAP generated using StabMap coloured by simulated data type (left), and by cell type (right). **e**. local inverse Simpson indices (LISI) for simulated data type (top row) and for cell type (bottom row). Each boxplot corresponds to different choices of number of cells in the multiome dataset. The dotted line indicates approximately 1,000 cells, where LISI values appear to markedly shift from unfavourable to favourable integration.

### Spatial mapping of mouse chimera data using StabMap identifies differences in abundance along major anatomical axis

A distinct advantage of mosaic data integration is the ability to integrate datasets where distinct features have been probed. An additional advantage is that the joint embedding can be used to facilitate downstream analyses, including differential abundance testing across experimental groups. To demonstrate this, we explored embryonic day (E)8.5 single cell RNA-seq data from the mouse^1^, together with perturbation experiment data in the form of Brachyury (T) knockout T^-/-^/WT chimeras and control WT/WT chimeras collected at the same time point^17^. Chimeric embryos contain a mix of host (Wildtype (WT)) cells and injected cells that are labelled with td-Tomato; the injected cells in the control chimera are WT, while the injected cells in the T^-/-^/WT chimeras lack a functional copy of Brachyury (T)^17^. We also considered single cell resolution spatially resolved seqFISH data from a similar developmental timepoint^7^. For the scRNA-seq datasets we considered the union of highly variable genes, whilst for the seqFISH data we considered all 351 genes that were probed in the experiment. Additionally, for the seqFISH data, we extracted new features, corresponding to the mean expression of each gene among the immediate neighbours of each cell, thus providing information about each cell’s local, spatially-resolved, context (Figure 4A, Methods). We used StabMap to jointly embed these data into the same latent space, and used fastMNN^18^ to correct for any batch effects among the individual pools for each experimental platform. We observed that all cell types separated well, with good mixing between data collected from each modality (Figure 4B).

**Figure 4.**
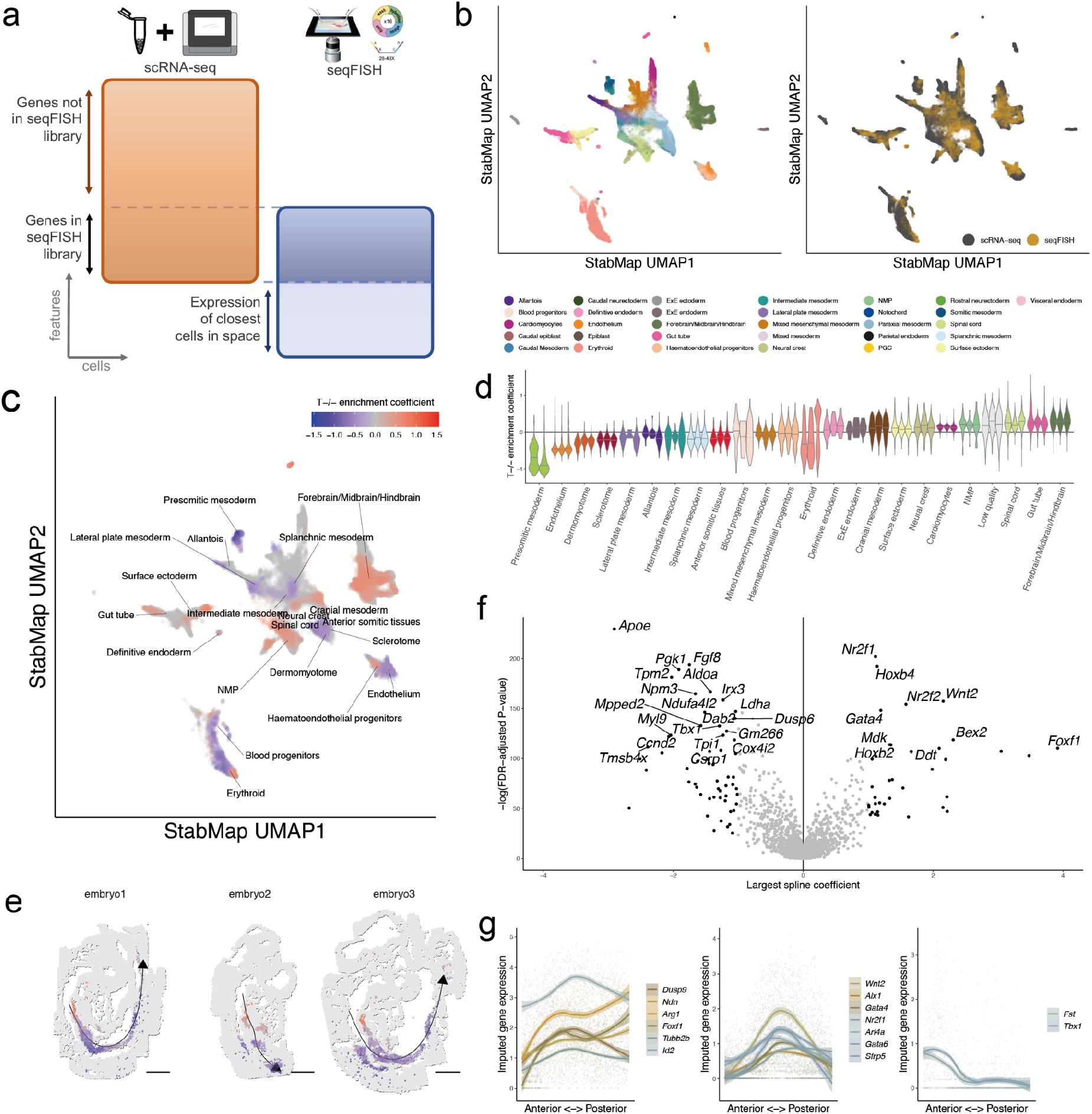
Integration of T-chimera and seqFISH data using StabMap with spatial neighbour feature extraction. **a**. Summary of mosaic data integration task and features used. Cells captured using scRNA-seq belonging to the E8.5 mouse gastrulation atlas ^1^, WT/WT chimera ^1^, and T^-/-^/WT chimera ^17^. seqFISH cells are obtained from sagittal sections of three E8.5 embryos [Citation error]. Features used for the scRNA-seq data are the union of the highly variable genes for each dataset. Features used for the seqFISH data are the gene expression of each cell, as well as the mean gene expression of the most proximal cells in space. **b**. UMAP plots displaying all cells after performing StabMap. Cells are coloured by the cell type (left) and by the platform (right). **c**. UMAP plot of all seqFISH cells coloured by local enrichment coefficient value of T^-/-^ enrichment test for statistically significant tests. **d**. Violin plots of T^-/-^ enrichment coefficients per embryo split by cell type. **e**. Spatial graphs of seqFISH embryos, with cells coloured by T^-/-^ coefficients for cells assigned a splanchnic mesoderm identity. Curved lines are fitted principal curves associated with the Anterior-to-Posterior (AP) axis along each embryo. **f**. Volcano plot showing value of largest magnitude spline coefficient (x-axis) and -log(FDR-adjusted P-value) for statistical test of splines model for splanchnic mesoderm (Methods). Highly ranked genes with large spline coefficients are labelled. **g**. Scatterplots and local mean expression ribbons of clustered genes showing distinct patterns of expression along the AP axis in splanchnic mesoderm.

Given this joint embedding, we next performed spatially-resolved enrichment testing of the relative abundance of T^-/-^ cells across the common space, to discover whether there are regions within the embryo where the T^-/-^ cells are enriched or depleted - an analysis that is only possible possible with the StabMap embedding. To do this, we first identified, for each seqFISH cell in the joint embedding, the 1,000 nearest neighbour cells from the T^-/-^/WT and the control WT/WT chimera samples. Among these 1,000 nearest neighbour cells, we calculated the relative fraction of cells contributing to the td-tomato+ population for each biological replicate of the T^-/-^ /WT and WT/WT samples. Subsequently, for each seqFISH cell, we used logistic regression to statistically assess whether there was a local enrichment or depletion of T^-/-^ cells (Methods), identifying 16,677 significant seqFISH cells (FDR-adjusted P-values < 0.05 out of a total of 57,536 seqFISH cells) (Figure 4C, Supplementary Figure 3A).

Upon examining the annotation of these cells, we found, consistent with previous analysis^17^, broad depletion of T^-/-^ cells among the Presomitic mesoderm, Dermomyotome, and Sclerotome alongside broad enrichment in neuromesodermal progenitors (NMPs) (Figure 4D, Supplementary Figure 3B). Intriguingly, we observed a heterogeneous distribution of local T^-/-^ enrichment in the splanchnic/pharyngeal mesoderm (42 cells displaying significant positive enrichment and 543 cells displaying significant negative enrichment (FDR-adjusted P-value < 0.05)), a cell type associated with tissues surrounding the forming gut. When we examined the physical locations of these cells, we observed an extremely strong concordance between the local T^-/-^ enrichment coefficient and the relative positioning of the cells along the anterior-to-posterior (AP) axis, as quantified using principal curves^19^ (Spearman correlation ranging between -0.26 and -0.68, Figure 4E, Methods).

We then used non-parametric cubic splines to identify imputed gene expression patterns that varied significantly along the principal curve (Figure 4F, Methods), and identified *Tbx1* and *Fgf8*, key genes regulating the development of anterior splanchnic mesoderm^20^ in the domain enriched for *T*^-/-^ cells. Conversely, markers of gut-associated splanchnic mesoderm *Foxf1 and Wnt2* (Figure 4G)^21,22^, and of posterior mesoderm homeobox genes *Hoxb2* and *Hoxb4* (Supplementary Figure 4) were enriched in the more posterior regions depleted in *T*^-/-^ cells.

Together, these observations suggest a broader role of Brachyury on regulating formation of posterior mesodermal tissues well beyond somitogenesis. In particular, this suggests that distinct domains of splanchnic mesoderm may also have distinct levels of dependency on Brachyury.

Critically, our spatial mapping of the relative enrichment of T^-/-^ cells using StabMap provides a basis for mapping complex experimental data onto a spatial reference, thereby allowing us to draw these inferences without the need to perform time-consuming and costly spatial perturbation experiments.

## DISCUSSION

In this paper, we have introduced StabMap, an approach to perform mosaic data integration for single cell data. StabMap accurately embeds single cell data from multiple technology sources into the same low dimensional coordinate space, using labelled or unlabelled single cell data, and performs well even when some dataset pairs do not share any features.

A current limitation of StabMap is that all features from an experiment are considered together. However, for single cell multiomics data an alternative would be to consider the different omics layers as individual data matrices, rather than to concatenate them into a large matrix^6^. This concatenation step corresponds to a naive example of vertical integration, where techniques such as feature standardisation are employed to ensure comparability across different modalities measured in the same cell. StabMap could be extended to employ more sophisticated vertical integration techniques, for example incorporating factors that describe variability across multiple layers, as implemented within MOFA^23^ or sharing information across multiple layers, as implemented within the weighted-nearest-neighbours framework^24^.

A key advantage of StabMap is the ability to incorporate novel analytical features, which may only exist for a subset of datasets, in the data integration step. We have demonstrated this using the spatial seqFISH data integration by using the expression of each gene in the most proximal cells in physical space as a feature (something that can not be captured in dissociated scRNA-seq data). Additionally, other bespoke features can be considered, such as local variance or local correlation values on spatial or trajectory-based data^25^, or cell-specific information such as lineage or clonal tracking information^26^. The ability to integrate data from such diverse sources offers the potential to extract novel biological insights by taking full advantage of diverse input datasets.

We envisage StabMap being used in a variety of contexts, especially as large-scale analysis of publicly available (and typically inconsistently processed datasets) becomes more widespread. Matching features between various datasets and ensuring a common data pre-processing pipeline is a serious hindrance for standard integration tools and can hinder the ability to draw biological insight. Consequently, StabMap could be employed to ensure that informative features are not lost purely due to practical challenges in pre-processing, enabling more comprehensive and complete downstream analysis.

## METHODS

### Mosaic data topology

The input to StabMap is a set of *s* appropriately scaled and normalised data matrices, 𝒟 = {*D*_1_, *D*_2_ … *D*_*s*_}, not necessarily containing the same features, and optional discrete cell labels for any of the datasets. As an initial step, StabMap generates the corresponding mosaic data topology (MDT). The MDT is an undirected weighted network which contains s nodes, one corresponding to each data matrix, with edges being drawn between pairs of nodes for which the corresponding data matrices share at least one feature. The edges in the MDT are weighted according to the absolute number of common features between the two datasets. StabMap requires that the MDT be a connected network, i.e. that there exists a path between any two nodes. Weighted shortest paths are calculated between any two given nodes in the MDT.

### The StabMap algorithm

At least one dataset must be considered as a reference dataset, with the option for multiple datasets to be considered as reference datasets. The output of StabMap is a common low-dimensional embedding with rows corresponding to all cells across all datasets, and columns corresponding to the sum of lower dimensions across the reference dataset(s). For a reference dataset *D*_*r*_, two matrices are extracted, first a scores matrix *S*_*r*_ (a cells x low-dimensions matrix) and a loadings matrix *A*_*r*_ (a features x low-dimensions matrix) such that 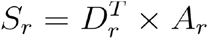. If no cell labels are provided, principal components analysis (default 50 PCs) is used for estimation of *S*_*r*_ (as the PC scores) and *A*_*r*_ (the components loadings). Alternatively, if discrete cell labels are provided, linear discriminant analysis is used for estimation of *S*_*r*_ (as the linear discriminants for each class) and *A*_*r*_ (the feature discriminant loadings).

Then, for each of the *s* data matrices, score matrices 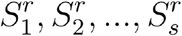 are calculated in one of the following ways for data matrix *i*:

- If *i* = *r*, then the scores matrix *S*_*r*_ is returned, i.e. 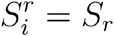;
- If *i* and *r* share an edge in the MDT, and all features in *A*_*r*_ are present in *D*_*i*_, then 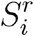 is directly calculated as the projected scores, i.e. 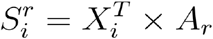, where *X*_*i*_ is the appropriate submatrix of *D*_*i*_ to match the features in *A*_*r*_. If not all of the features in *A*_*r*_ are present in *D*_*i*_, then 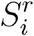 is estimated using multivariate linear regression on each column of *S*_*r*_ for dataset *D*_*r*_. Specifically, for column *j* of *S*_*r*_, we fit the model *S*_*r*_ [*j*] = *X*_<*r,i>*_[*j*] *β*_<*r,i>*_[*j*] + *ϵ* where *X*_<*r,i>*_ is the submatrix of *D*_*r*_ for features that are shared among *D*_*i*_ and *D*_*r*_, and *ϵ* is assumed to be normally distributed noise. *B*_<*r,i>*_ therefore is a matrix of fitted coefficients 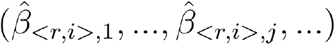 with rows corresponding to the shared features between *D*_*i*_ and *D*_*r*_ and columns corresponding to the columns of *S*_*r*_. The estimated score matrix for *i* is taken to be the predicted values of the multivariable linear model for dataset *D*_*i*_, and is calculated as 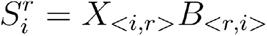 where *X*_<*i,r>*_ is the submatrix of *D*_*i*_ for features that are shared among *D*_*i*_ and *D*_*r*_.
- If *i* and *r* do not share an edge in the MDT, then 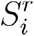 is estimated using an iterative approach that exploits the shortest weighted path in the MDT. Starting from node *r*, for the next node along the path *p*, we calculate 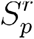 as described above. If the next node along the path is *i*, then we fit the model 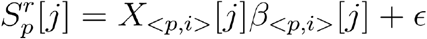 where *X*_<*p,i>*_ is the submatrix of *D*_*p*_ for features that are shared among *D*_*p*_ and *D*_*i*_ and *B*_<*p,i>*_ is the matrix of fitted coefficients 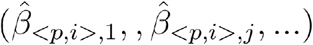. The estimated score matrix for *i* is then taken as the predicted values of this multivariable linear model for dataset *D*_*i*_, and is calculated as 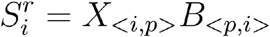. If instead, the next node along the path from *r* to *p* and eventually to *i* is some other node *q*, then this process of fitting a multivariable linear model and predicting on the new data is repeated until we calculate 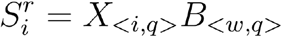, where *w* is the node previous to *q* along the path between *r* and *i*.

The estimated score matrices for each of the *s* datasets are then concatenated across rows to form the joint low dimensional score where reference *r* is employed: 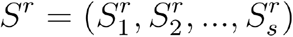, where *S*^*r*^ is a matrix with number of rows equal to the total number of cells across all *s* datasets and number of columns equal to the number of columns (selected features) in *S*_*r*_.

### StabMap with multiple reference datasets

For the set of reference datasets ℛ = {*D*_*j*_ s.t. *j* is in reference indices} ⊆ 𝒟, we calculate the corresponding set of joint low dimensional scores as described above, 𝒮 = {*S*^*j*^ s.t. *j* is in reference indices}. We reweight each scores matrix *S*^*j*^ according to the overall L1 norm of the matrix and a user-set weighting parameter *w*_*j*_ ∈ [0,1] (by default set to 1),

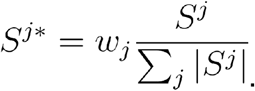

The user-set weighting parameter *w*_*j*_ controls the magnitude of the score vectors for each reference dataset, and thus corresponds to the relative influence of the reference dataset on any magnitude-based downstream analysis (e.g. calculation of Euclidean distances between cells). To generate common low dimensional scores across all reference datasets, we concatenate the re-weighted scores across columns to form the StabMap low dimensional scores, 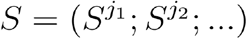 for reference data indices *j*_1_, *j*_2_, …. *S* is a matrix with number of rows equal to the total number of cells across all *s* datasets, and number of columns equal to the total number of columns across the scores matrix for each reference dataset.

### Downstream analysis with StabMap

#### Batch correction

While StabMap jointly embeds cells across multiple datasets into a common low dimensional space, batch effects both within and among datasets can remain. Any existing batch correction algorithm that works on a low dimensional matrix (e.g. fastMNN^18^, scMerge^27^, BBKNN^28^) can be employed to obtain batch-corrected StabMap embeddings. In the analyses presented in this manuscript we use fastMNN but note that users are able to apply any suitable algorithm for this task.

#### Supervised and unsupervised learning

The batch-corrected StabMap embedding facilitates supervised learning tasks such as classification of discrete cell labels using any suitable method such as k-nearest neighbours, random forest, and support vector machines, and regression using traditional linear models or support vector regression. Unsupervised learning tasks can be performed by clustering directly on the embedding (e.g. k-means clustering) or by first estimating a cell-cell graph (e.g. shared nearest neighbour or k-nearest neighbour graph) followed by graph-based clustering (e.g. louvain or leiden graph clustering). Since one can use the embedding to estimate the cell-cell graph, additional bespoke single cell analyses such as local differential abundance testing between experimental groups, such as that implemented in Milo^29^ can be employed.

#### Imputation of original features

We include a imputation implementation based on the StabMap low-dimensional embeddings to predict the full-feature matrices for all data, by extracting the set of k neighbours using Euclidean distance within the StabMap-projected space, and returning the mean among the nearest neighbours. This is especially useful for projecting query data onto a reference space or for identifying informative features downstream of the data integration step.

### Mosaic data integration simulations

We used publicly available data to investigate the performance of StabMap and other methods, as described below.

#### PBMC 10X Multiome data

We used the SingleCellMultiModal R/Bioconductor package^30^ to download the ‘pbmc_10x’ dataset, containing gene expression counts matrix and read counts associated with chromatin peaks captured in the same set of cells. We normalised the gene expression values using logNormCounts^31^ in the scuttle package, and restricted further analysis to highly variable genes (HVGs) selected using the ModelGeneVar function in scran^32^. For the chromatin data modality we performed term frequency - inverse document frequency (TF-IDF) normalisation according to the method described in^10^. We extracted peak annotation information using the MOFA2 R package tutorial^23^, including information on which genes’ promoters the chromatin peaks were associated with, if any. These promoter peaks were annotated as the associated gene name, so that the promoter peak features would match the RNA genes features.

To perform the mosaic data integration simulation with the PBMC 10X Multiome data, we ignored the matched structure between the RNA and chromatin modalities, and treated this data as if it belonged to two distinct datasets. We performed StabMap using both RNA and chromatin modalities as the reference datasets, and re-weighted the embedding to give equal contribution for the two modalities. For assessing the cell type accuracy we used the RNA modality cells as labelled data, and predicted the cell types of the chromatin modality cells using k-nearest neighbours classification with k = 5.

#### Mouse Gastrulation Atlas scRNA-seq

We downloaded the counts data from Pijuan Sala et al (2019) using the MouseGastrulationData R/Bioconductor package^33^ corresponding to embryonic day (E)8.5, and normalised and extracted HVGs in the same way as the 10X Multiome PBMC data. For the simulation, we randomly selected 100, 200, 500, and 1000 genes from among the HVGs. Then, we split the dataset into four groups according to the four sequencing samples. For each randomly selected pair of sequencing samples, we artificially assigned one sequencing sample as the query dataset, restricting to the randomly selected genes, and kept one other sequencing sample intact as the reference dataset.

We used StabMap to jointly embed the reference and query datasets into a common low dimensional space by selecting the reference dataset as the sole reference, followed by batch-correction using fastMNN. We also performed naive PCA, UINMF and MultiMAP for comparison. To assess performance, we calculated the mean accuracy of cell type classification of query cells using k-Nearest Neighbours with k = 5 for each method.

#### Comparison with other methods

##### UINMF

We used software version 0.5.0 of LIGER, which includes the UINMF implementation, and performed integration using defaults as suggested in the LIGER vignette. We used the counts matrix for input, as suggested in the vignette. We used the resulting 50-dimensional embedding for subsequent downstream analysis, and UMAP implemented in scater^31^ for visualisation.

##### MultiMAP

We used the Python (version 3.8.10) package MultiMAP (version 0.0.1), and performed data integration using defaults as suggested by the MultiMAP tutorial website with equal weights for each dataset. The output of MultiMAP is a corrected graph representation, as well as a two-dimensional representation of the data. We used this two-dimensional representation for visualisation and to perform downstream analysis tasks.

##### Naive PCA

To implement naive PCA, we first extracted the submatrices of datasets containing features that were common across all datasets. We then performed PCA using scran’s implementation with 50 principal components, followed by batch correction using MNN. We used the 50-dimensional representation for downstream analysis tasks, and UMAP to perform further dimensionality reduction to two dimensions for visualisation.

#### Evaluation

To evaluate the mosaic data integration simulations, we used three quantitative metrics.

##### Cell type classification accuracy

Given a joint embedding, we perform a simulation such that discrete class labels corresponding to cell types are artificially removed for a subset of the data. We then perform K-nearest neighbours classification (k = 5) to obtain the predicted class label for the artificially unlabelled data. The cell type classification accuracy is thus the proportion of cells for which the classification is correct compared to the true cell type label,

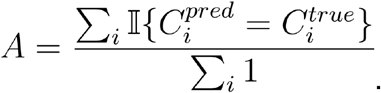

##### Jaccard similarity

For cell *i* in embedding *S* we have *l* positions for the *l* omics levels (e.g. RNA, chromatin). We extract the sets of size *k* (default 100) containing the nearest cells of the same omics layer, i.e.

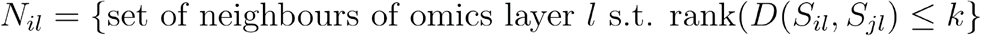

where *D* (*a,b*) is the Euclidean distance of vectors *a* and *b*. The Jaccard similarity is thus

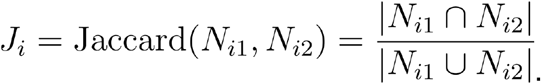

Larger values of *J*_*i*_ correspond to larger overlap of neighbours between the two omics layers and are thus desired.

##### Number of nearest cells metric

Similar to the metric employed by Kriebel et al. and Jain et al. ^11,12^, for cell *i* belonging to omics layer 1 (e.g. RNA) in embedding *S*, we calculate the number of cells among omics layer 2 (e.g. chromatin) which are nearer than cell *i* belonging to omics layer 2, 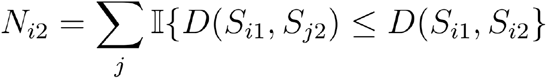.

We then extract the empirical cumulative distribution of nearest cells by calculating, for each integer *x*, the number of cells for which their number of nearest cells metric is at most this value, 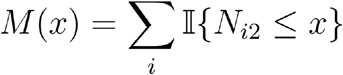. Higher values of *M*(*x*) across all values of *x* are more desired.

### Disjoint mosaic data integration simulation

We used the PBMC 10X Multiome data to evaluate StabMap under the situation of disjoint mosaic data integration. We downloaded and processed the data as described in the subsection above, with the exception that promoter peaks corresponding to specific genes were not matched to the associated genes. This resulted in a complete lack of overlap between features between the RNA and chromatin modalities.

To perform the simulation, we randomly allocated each cell into one of three classes: 1) RNA only, 2) chromatin only, and 3) Multiome, with varying relative proportions of cells associated with the Multiome class. Cells within the RNA class had their chromatin information ignored, and cells within the chromatin class had their RNA information ignored, while cells within the Multiome class were left unchanged. We then used StabMap to integrate these three simulated datasets and generate a low-dimensional embedding for each simulation setting. Comparison with other methods is not possible since PCA, UINMF and MultiMAP require at least some overlapping features across all datasets.

To evaluate the disjoint mosaic data integration simulation, we calculated the local inverse Simpson index (LISI)^16^ using both modality and cell type as the grouping variables. Higher LISI values correspond to more local mixing of cells, and so relatively high values for modality and low values for cell type are desirable.

### Spatial mapping of mouse chimera data using StabMap

#### scRNA-seq data

We used the MouseGastrulationData R/Bioconductor package (Griffiths and Lun 2020) to download gene expression counts for the Mouse Gastrulation Atlas dataset, WT/WT control chimera dataset^1^, and T^-/-^/WT chimera dataset^17^, corresponding to embryonic day (E)8.5. We combined the gene expression counts into a single dataset, then normalised and extracted HVGs using the same approach applied to the 10X Multiome PBMC data.

#### seqFISH data

We downloaded seqFISH-resolved gene expression log-counts^7^ for spatially-resolved cells of mouse embryos profiled at a similar developmental stage along with their corresponding spatial coordinates. We extracted novel features for each gene *g* and each cell *i* by calculating the mean expression value among the nearest cells in space,

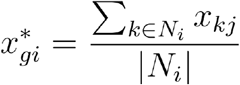

where *N*_*i*_ = {*k* s.t. *D*(*i,k*) ≤ 2,*i* ≠ *k*} is the set of cells that are at most 2 steps away from cell *i* in the spatial nearest neighbour network^7^. We then concatenated these novel features with the measured gene expression, prior to downstream integration with the dissociated scRNA-seq data.

#### Mosaic data integration and local enrichment testing

We used StabMap, parametrised with multiple reference datasets, to integrate the scRNA-seq and seqFISH data. We used PCA (default 50 PCs) to generate the low dimensional scores for the scRNA-seq and seqFISH references, and reweighted each scores matrix using the default weighting parameter of 1. As a result, we obtained a 100-dimensional StabMap low dimensional scores matrix. We then corrected for any remaining batch differences using fastMNN, where batches reflect technical groups from each dataset.

To calculate whether T^-/-^ cells were enriched in a neighbourhood around each seqFISH cell, we performed logistic regression. Specifically, for each spatially-resolved (seqFISH) cell, in the joint embedding we extracted its 1,000 nearest neighbours from each chimera dataset (4 T^-/-^/WT samples and 3 WT/WT samples, and fit the model 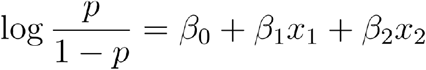.

In this model, *p* is the vector of observed proportions of td-tomato+ cells for each chimera, *x*_1_ is a vector containing the total proportion of td-tomato+ cells belonging to a biological replicate, and *x*_2_ is a vector indicating whether a chimera is T^-/-^/WT or WT/WT. We extracted the estimated coefficient of interest, 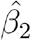, and associated P-value for each spatially resolved cell using a likelihood ratio test, resulting in a local measure of enrichment or depletion of T^-/-^ cells for each seqFISH-profiled cell. We then used the method of Benjamini-Hochberg to calculate FDR-adjusted P-values.

#### Mixed T^-/-^ enrichment in pharyngeal/splanchnic mesoderm

To examine the relationship between the estimated T^-/-^ enrichment coefficient and anterior-posterior (AP) axis position in the splanchnic mesoderm, we fitted principal curve models, with 4 degrees of freedom, for each individual spatially resolved embryo with the spatial coordinates as the underlying data^19^. We used the principal curve fitted values to extract the AP ranking of cells along this axis, and then used this ranking to estimate a locally smoothed T^-/-^ enrichment coefficient along the AP axis.

To assess gene expression changes along the AP axis as T^-/-^ cells move from being enriched to being depleted, we selected an equal number of cells anterior and posterior to the position where the smoothed T^-/-^ enrichment coefficient is zero, and performed differential gene expression analysis using imputed gene expression values. Imputed gene expression was quantified for each spatially-resolved cell using the mean gene expression value of the nearest five Mouse Gastrulation Atlas cells in the StabMap low-dimensional space. Gene expression changes along the AP axis were assessed using a non-parametric cubic splines model with 3 degrees of freedom along with grouping variables for the individual embryos. Statistical significance was estimated using an F-test, with a null model of no splines effects, with empirical Bayes shrinkage using the limma framework, followed by adjustment for multiple testing. For statistically significant genes, we visualised gene expression along the AP axis using local loess smoothing and ribbon plotting for the local standard error.

## SUPPLEMENTARY FIGURE LEGENDS

**Supplementary Figure 1.**
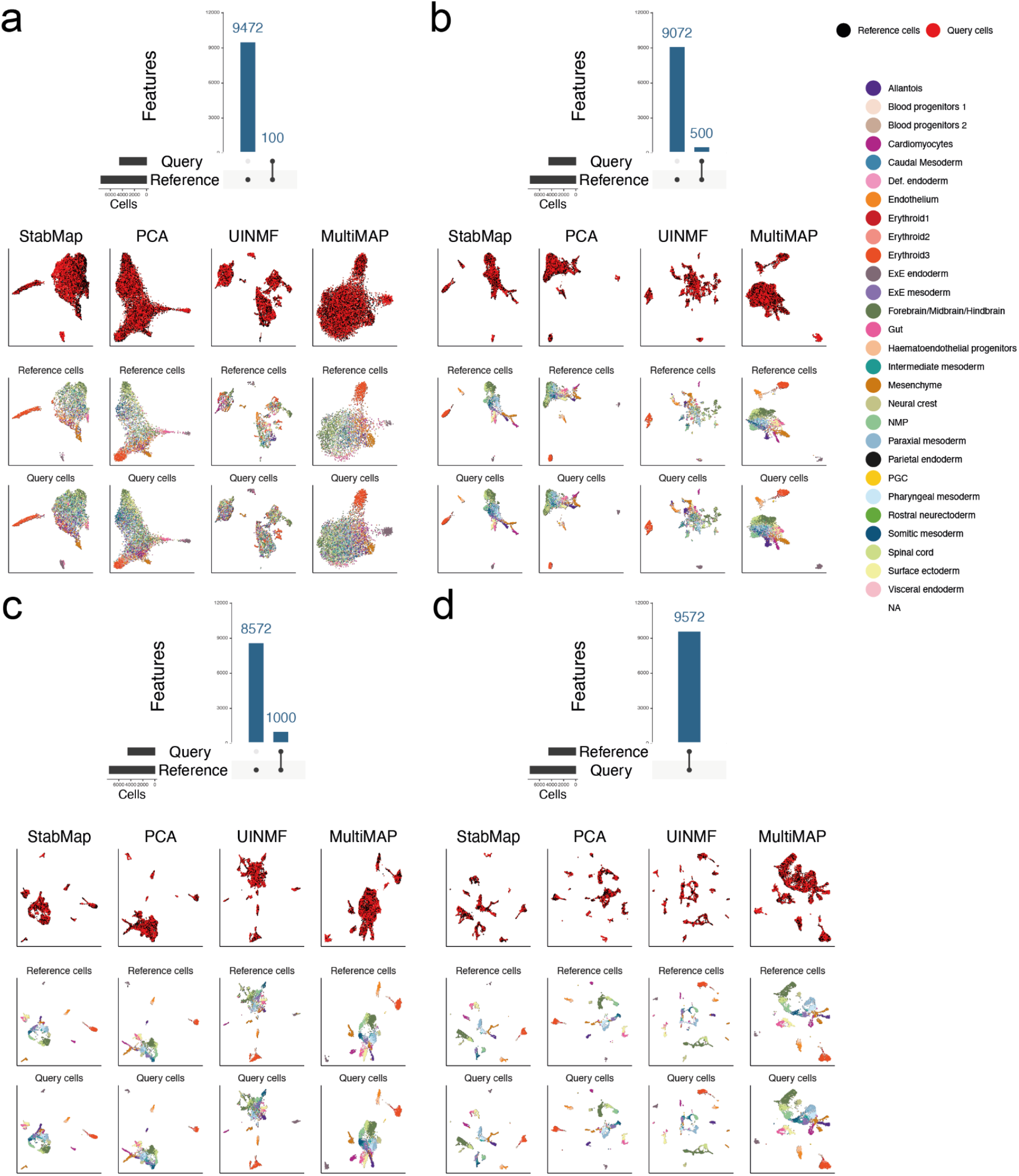
**a**. UpSet plot and UMAP representations of Mouse Gastrulation Atlas data simulation with 100 randomly selected features using StabMap, PCA, MultiMAP, and UINMF. First row shows the query cells coloured by simulated dataset, the second row shows reference cells coloured by cell type, and the third row shows query cells coloured by cell type. **b-d**. As in panel (a.) for 500, 1,000, randomly selected and all features respectively.

**Supplementary Figure 2.**
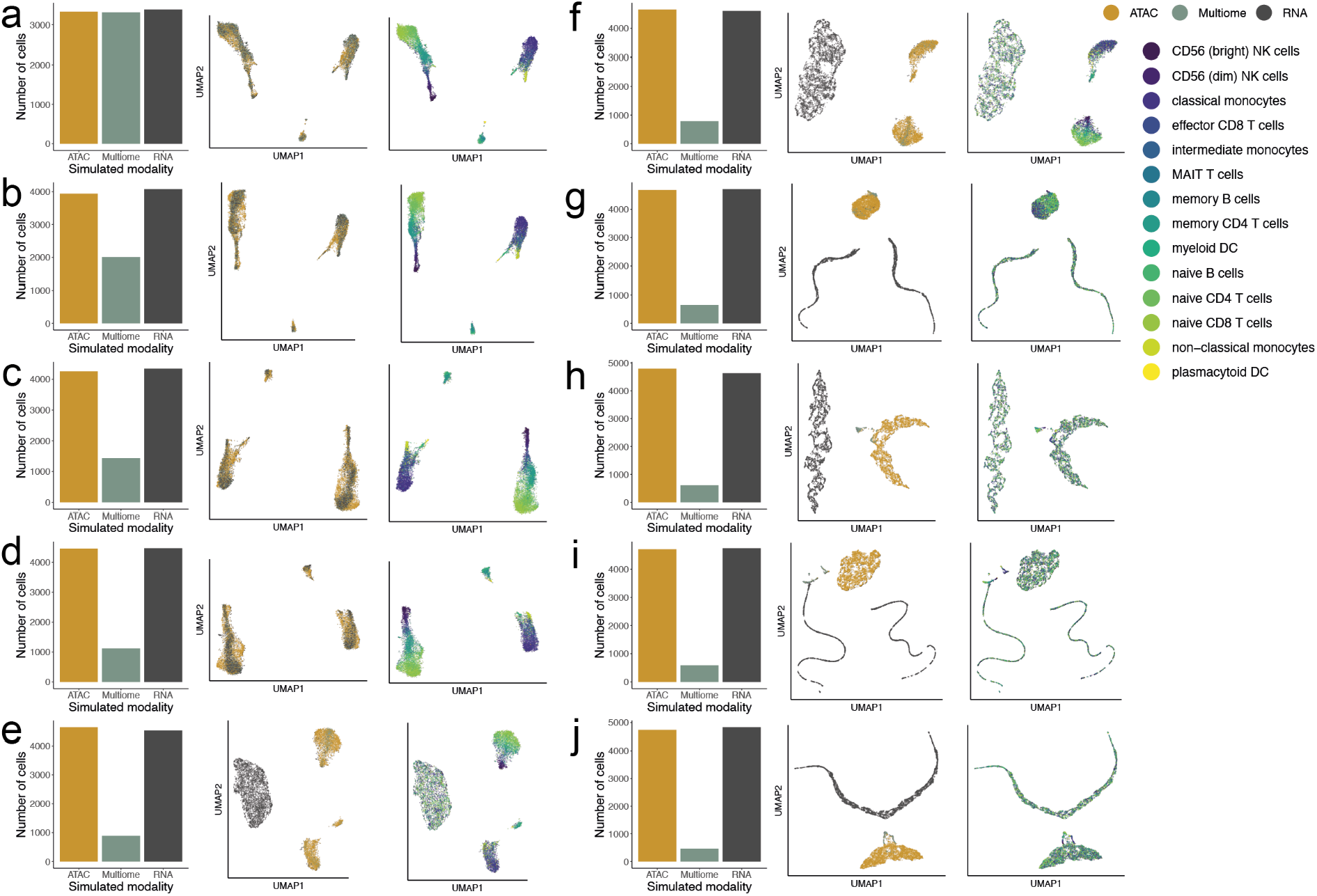
**a**. Number of 10X PBMC Multiome cells assigned to each simulated data type (left), joint UMAP generated using StabMap coloured by simulated data type (middle), and by cell type (right). **b-j**. As in panel (a.) for decreasing proportions of simulated Multiome cells.

**Supplementary Figure 3.**
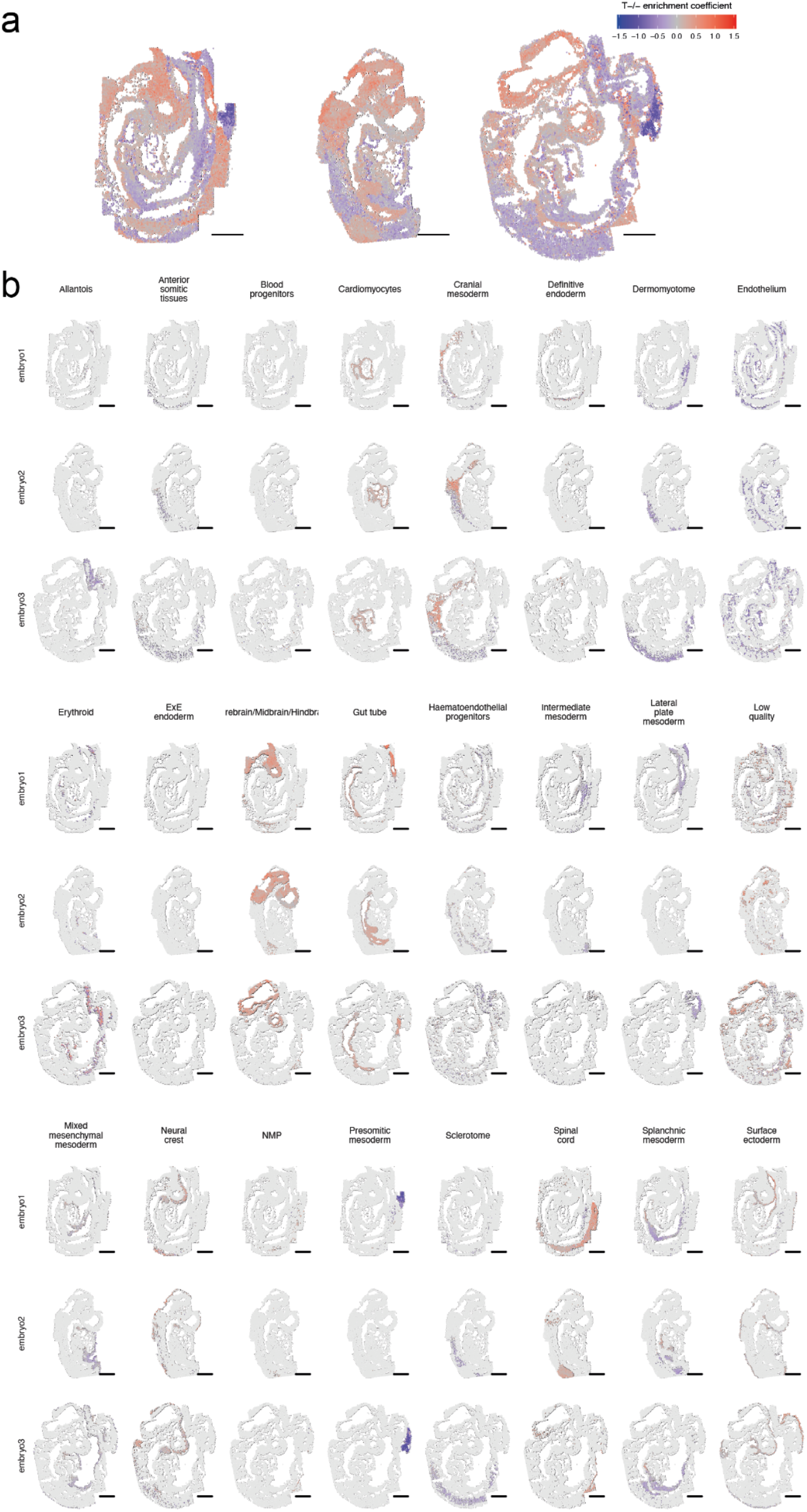
**a**. Spatial coordinates plot of all seqFISH cells coloured by local coefficient value of T^-/-^ enrichment test. **b**. Spatial coordinates plots of all seqFISH cells, split by cell type (columns) and embryos (rows), where selected cells are coloured by local coefficient value of T^-/-^ enrichment test.

**Supplementary Figure 4.**
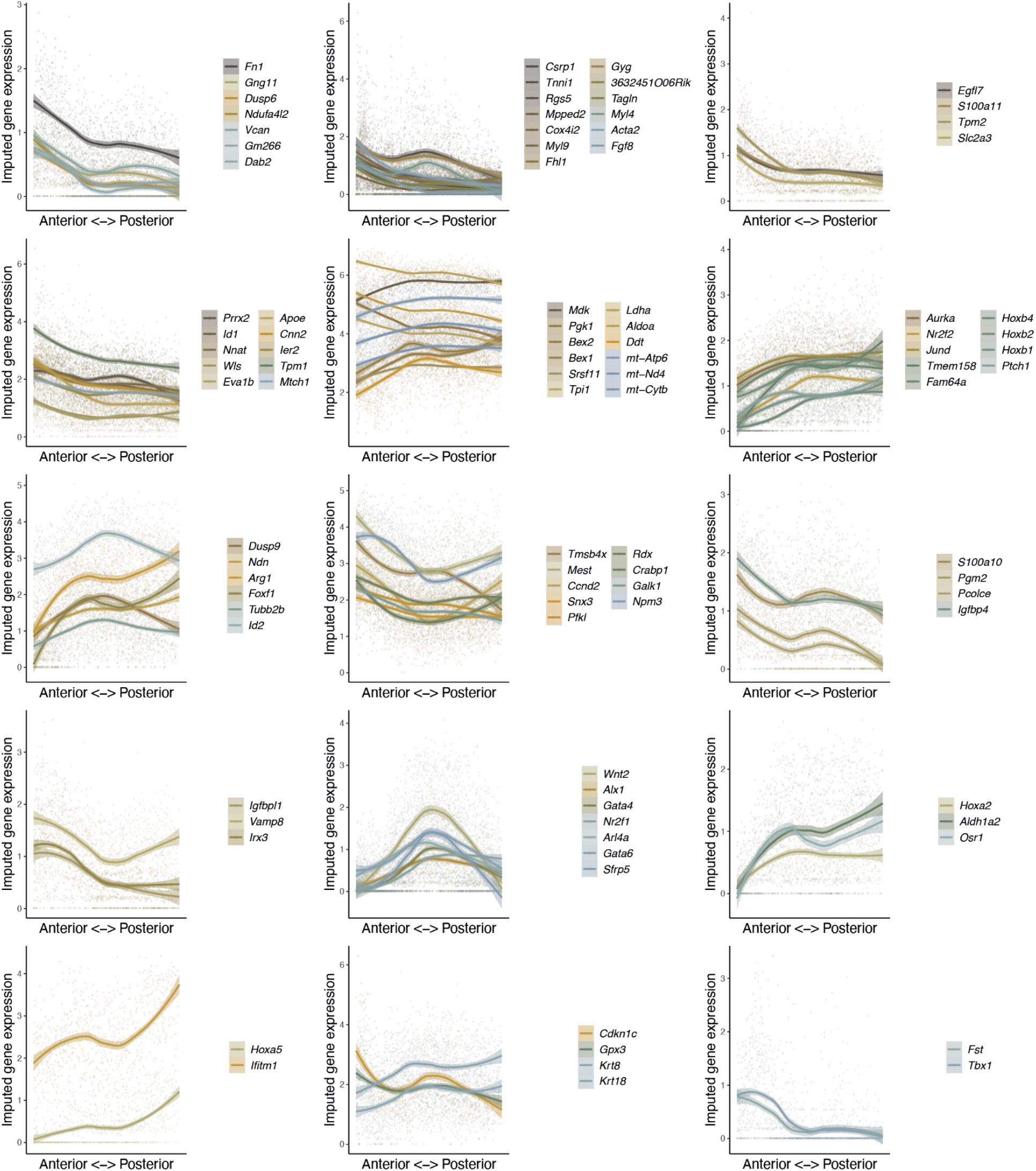
Scatterplots and local mean expression ribbons of significantly varying genes (cubic splines likelihood ratio test FDR-adjusted P-values < 0.05), clustered using hierarchical clustering to show distinct patterns of expression along the AP axis in splanchnic mesoderm.

## DATA AVAILABILITY

This study used publicly available data. The PBMC 10X Multiome and mouse embryo scRNA-seq data were accessed via Bioconductor (version 3.13) ExperimentHub packages MouseGastrulationData (version 1.6.0) and SingleCellMultiModal (version 1.4.0) respectively. The processed mouse embryo seqFISH data was accessed online via the web portal https://marionilab.cruk.cam.ac.uk/SpatialMouseAtlas/.

## CODE AVAILABILITY

All analyses were performed in R (version 4.1.0). The StabMap software is available as an R package at https://github.com/MarioniLab/StabMap. Scripts for analysis and figure panels in this manuscript are available at https://github.com/MarioniLab/StabMap2021.

## ACKNOWLEDGEMENTS

We thank our colleagues in the University of Cambridge Cancer Research UK Cambridge Institute, University of Gothenburg, and the EMBL European Bioinformatics Institute for their support and intellectual engagement. We thank M. Morgan, D. Keitley, A. Missarova, E. Heidari and other members of the Marioni lab for discussions concerning the analysis. We thank M-L. Ton, T. Lohoff, and R. Argelaguet for their discussions surrounding the project.

## AUTHOR CONTRIBUTIONS

S.G. and J.C.M. conceived the study. S.G. developed the method and software and performed data analysis with input from J.C.M. S.G. interpreted the results with input from C.G. and J.C.M. S.G., J.C.M. and C.G. wrote the manuscript. All authors read and approved the final version of the manuscript.

